# Directional and frequency characteristics of auditory neurons in Culex male mosquitoes

**DOI:** 10.1101/608778

**Authors:** Dmitry N. Lapshin, Dmitry D. Vorontsov

## Abstract

The paired auditory organ of mosquito, the Johnston’s organ (JO), being the receiver of particle velocity component of sound, is directional by its structure. However, to date almost no direct physiological measurements of its directionality was done. In addition, the recent finding on the grouping of the JO auditory neurons into the antiphase pairs demanded confirmation by different methods. Using the vector superposition of the signals produced by two orthogonally oriented speakers, we measured the directional characteristics of individual units as well as their relations in physiologically distinguishable groups – pairs or triplets. The feedback stimulation method allowed to discriminate responses of the two simultaneously recorded units, and to show that they indeed responded in antiphase. We also show that ratios between the individual tuning frequencies in pairs and triplets are non-random and follow the principle of harmonic synchronization, remarkably similar to the one known from the observations of mosquito behavior. Units of different tuning and sensitivity are evenly distributed around the axis of the JO, providing the mosquito with the ability to produce complex auditory behaviors.

**Summary statement:** Auditory neurons of mosquito are grouped into pairs or triplets, each unit tuned to a specific frequency. Within the pair units respond to opposite directions of the sound. Units of different tuning and sensitivity are evenly distributed around the axis of the Johnston’s organ.

## 1. Introduction

The ear of mosquito, the Johnston’s organ (JO), is a highly sophisticated system containing extremely large number of sensory neurons, measured in thousands (Boo and Richards, 1975a; Boo and Richards, 1975b; Hart et al., 2011). Its complexity should not be surprising since mosquito mating behavior depends on audition and hence the ears were developed under high selection pressure. Morphologically, mosquito possesses two feather-like antennae, designed to be the receivers of air particle velocity. The antenna originates from the capsule with the radially arranged sensillae, or scolopidia, most of which contain two or three bipolar sensory neurons. The neurons respond to antenna vibrations by transducing them into electrical potentials and send the axons to the brain via the antennal nerve.

The task of reception of a mate’s flight tone which lies within the narrow frequency range does not seem to be too complicated at the first sight. However, the real-life task which is solved by the mosquito auditory system is much more difficult. First of all, any external sound blends with mosquito’s own flight tone, that leads to the appearance of multiple mixed harmonics at the receptor input (Gibson et al., 2010; Lapshin, 2011; Lapshin, 2012; Simões et al., 2016; Simões et al., 2018; Warren et al., 2009). The flight tone itself is not stable since it depends on the ambient temperature (Sotavalta, 1952; Villarreal et al., 2017), and, in addition to this, mosquitoes continuously maneuver, change their flight velocity and, hence, the wingbeat frequency. This change is especially remarkable during the courtship ‘acoustic dance’, when male mosquitoes first produce the rapid frequency modulation, and then the pair of mosquitoes mutually tune their wingbeats to fit the specific frequency ratio (Aldersley and Cator, 2019; Cator et al., 2009; Gibson and Russell, 2006; Pennetier et al., 2010; Warren et al., 2009). A male mosquito not only needs to detect the presence of a female, but has to locate her and follow until copulation. And, since many mosquitoes mate in a swarm, to win the competition our male must perform these tasks faster than other males.

Given all this, the complexity of the mosquito JO is no more surprising. The principles of its operation are still waiting to be understood, and there is no reason to consider them being trivial. From the obvious tasks mentioned above one can presume that the JO must contain units tuned to different frequencies and, most probably, with different sensitivity to provide high dynamic range. The radially symmetrical flagellar JO is said to be inherently directional (Belton, 1974; Robert, 2005). At the level of sensory neurons this means that there must be sensory units selectively responding to a sound coming from any angular range, but whether the sensitivity and frequency preference of the JO is also symmetrical has never been tested, except several experiments made by Belton (1974).

In the much simpler organized JO of Drosophila (>400 mechanosensory neurons), different types of primary sensory neurons were discovered (Albert and Göpfert, 2015; Kamikouchi et al., 2009; Yorozu et al., 2009), as well as interneurons which selectively code the complex stimulus features (Chang et al., 2016; Matsuo and Kamikouchi, 2013). Although the Drosophila model allows to analyze the mechanisms which are not currently accessible in mosquitoes, the results of these studies should be transferred to the mosquito audition with some caution due to much higher complexity of the latter and substantially different acoustic behavior in fruit flies and mosquitoes. The physiological approach directed to testing the properties of the auditory neurons in mosquito involves the recording of their responses to sound. However, the method of recording the field potential of the whole JO, which is most commonly applied in the studies of the mosquito audition, does not allow to test any hypotheses on the diverse tuning of elements within the JO, neither frequency nor directional. Recently we developed a method of recording from small groups of axons of the JO sensory neurons along with controlled acoustic stimulation (Lapshin and Vorontsov, 2013). Although even the fine glass electrode used in that study allowed recording of focal potentials, still due to the extremely tiny diameter of the axons in the antennal nerve the resolution of recording had to be further improved, which was done by applying the positive feedback stimulation. Using it, it was possible to see the frequency preference of a unit situated closer to the electrode tip or possessing the lower threshold than other units.

The first finding which that study brought was the difference in frequency tuning between the JO sensory neurons both in female (Lapshin and Vorontsov, 2013) and male mosquitoes (Lapshin and Vorontsov, 2017). It was rather expected, keeping in mind the high number of units and also the behavioral observations which implied the frequency discrimination in mosquitoes (Aldersley and Cator, 2019; Aldersley et al., 2016; Lapshin and Vorontsov, 2018; Simões et al., 2016; Simões et al., 2018). The individual tuning frequencies were distributed from 85 to 470 Hz in males and from 40 to 240 Hz in females of *Culex pipiens*, while the majority of units was found to be tuned to the tones other than the wingbeat frequency of a mate.

A rather unexpected finding, however, was that the units going closely in the antennal nerve were found to be grouped in distinct pairs or triplets by the specifics of their response. In each group, the ratio between the individual tuning frequencies was non-random, while the frequencies themselves were distributed more or less randomly within the range of sensitivity. Moreover, the units within a pair demonstrated the antiphase response to the same stimulus. Each of the proposed explanations of the latter finding was nontrivial: either the axons from the opposite sectors of the JO were pairwise combined in a nerve, or the two units belonged to the same sensilla, but nonetheless responded in antiphase due to some specific tuning at the stage of mechanotransduction. The set of non-random ratios between the individual tuning frequencies within each pair or triplet of units remarkably coincided with the recent behavioral data on the mutual frequency tuning (Aldersley et al., 2016). Here we study the current larger set of measurement data in attempt to understand the principles behind the primary signal analysis which, as we presume, is performed by the JO sensory neurons even before this information enters the brain.

The weakness of our previous study was in the specifics of the positive feedback stimulation setup, which at that time provided only two variants of the stimulation phase: either 0° or 180° relative to the unit response. This limitation did not allow to tell whether the units which responded in a pair were strictly antiphase (180°), or possessed some other close directional ratio, since no directional measurements were performed. This was the primary reason for us to undertake another study and to utilize a different method of measuring the directional properties of the JO neurons. Here, we measure the thresholds of auditory neurons of the JO as a function of orientation of the acoustic wave vector relative to the axis of the antenna. Such data plotted in polar coordinates is commonly referred to as the polar pattern or the directional characteristic.

In previous studies, the polar pattern was assessed either by changing the angular position of the speaker relative to the test object (Daley and Camhi, 1988; Vedenina et al., 1998), or by rotating the test object relative to the stationary speakers (Hill and Boyan, 1976; Morley et al., 2012). There exists, however, a third way to study the directionality: to use a vector superposition of acoustic waves at the point of receiver, produced by two orthogonally oriented stationary speakers (Theunissen et al., 1996). In this study we implemented, with modifications, the latter method. It allowed us to avoid the mechanical vibrations associated with the movements of objects within the setup, that was in its turn important for stability of recording. Additionally, herein we compare the directional characteristics measured from the same sensory units by the sinusoidal and the positive feedback stimulation, and give the distribution of units with different frequency tuning around the axis of the mosquito antenna.

The preliminary study of directional properties of the JO sensory units was done in Chironomidae midges (Lapshin, 2015). Their auditory behavior is generally similar to that of Culicidae: swarming males are attracted to the female wingbeat tone; morphologically their JO’s also have many similarities. These experiments have shown that positive feedback stimulation can be successfully used to simultaneously measure the frequency and directional characteristics of the JO sensory neurons.

### 2. Methods

The relative threshold characteristics of auditory sensory units of the JO were measured depending on the orientation of acoustic wave vector relative to the axis of the antenna. In parallel, the individual tuning frequencies of units were identified.

### 2.1. Animal preparation

Males of *Culex pipiens pipiens* L. (n=91) were captured from a natural population in Moscow region of the Russian Federation. Experiments were conducted in laboratory conditions with air temperature 18-21°C in at the Kropotovo biological station (54° 51’ 2” N; 38° 20’ 58” E).

Individual mosquitoes were glued to a small (10×5 mm) copper-covered triangular plate by a flour paste with 0.15 M sodium chloride added. This type of attachment simultaneously serves three functions: it ensures good electrical contact of the mosquito with the plate, which was used as a reference electrode, mechanically fixes the mosquito and prevents it from drying during the experiment. The head of the mosquito was glued to its body by a bead of varnish to keep its orientation fixed during the experiment. The mosquito could still move its antennas, but this was visually controlled. The plate with the mosquito was mounted on a holder using a pair of miniature ferrite magnets which allowed to position the mosquito at any desired angle relative to the speakers. In most experiments the mosquito was positioned dorsal side up. However, the constant orientation of mosquito relative to the recording electrode could result in selective recording of some particular groups of neurons. To avoid this kind of a bias, the orientation was changed from specimen to specimen, either by turning it the ventral side up or by rotating it by 180° in the horizontal plane; measurements of directional responses were corrected accordingly. All recordings were made from the left JO.

### 2.2. Acoustic stimulation

Sound stimuli were delivered through a pair of Scandinavia 75 dynamic speakers (DLS, Sweden) positioned with their acoustic axes at the right angle (Fig. 1). The mosquito was fixed at the intersection of these axes in such a way that the base of the antenna flagellum and the axis of the associated JO was perpendicular to the plane defined by the axes of the two speakers.

**Fig 1.**
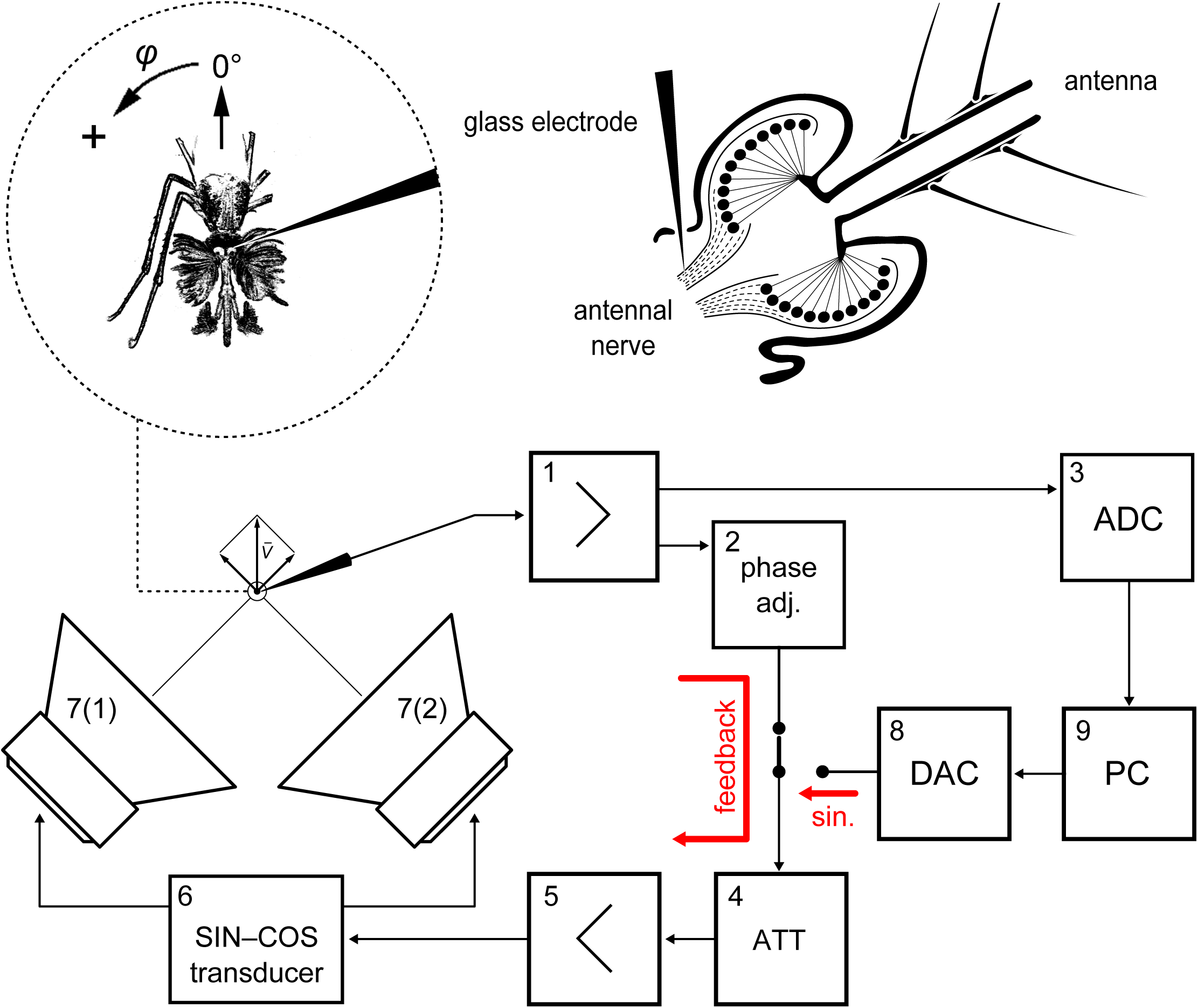
Experimental procedure. The experimental setup for electrophysiological recording and sound stimulation. Mosquito is fixed above two orthogonally oriented speakers. Neuronal response from the antennal nerve are amplified (1) and digitized (3) and stored on the PC (9). Sound stimulation is made alternatively in feedback mode (neuronal response after phase adjustment (2) via attenuator (4), power amplifier (5) and Sin-Cos transducer (6) is fed to speakers (7)) or sinusoidal mode (signal is synthesized on the PC (9) and after digital-to-analog converter (8) is fed to attenuator (4) and further to the speakers (7)). The mosquito is positioned at the intersection of axes of the two speakers in such a way that the base of the antenna flagellum is perpendicular to the both axes. The resulting direction of the air vibration velocity is determined by the vector superposition of the signals from the two speakers. An increase in angle of stimulation φ corresponds to counter-clockwise rotation of the velocity vector, with the insect’s head viewed from the front. Accordingly, when viewed from the mosquito’s head along the antenna the clockwise rotation corresponds to the increase in φ.

The speakers were powered from the home-made amplifier (K = 4) via a passive Sin–Cos (SC) transducer which produced two derived signals with the amplitudes,

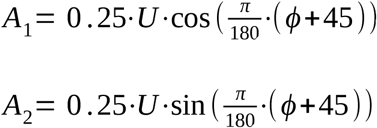

where A_1_ and A_2_ are the amplitudes of the control signals for the first and the second speaker, respectively; U is the alternating voltage at the input of the SC transducer; φ is the angle between the dorso-ventral line passing through the mosquito’s head and the vector of vibration velocity of air particles. An increase in φ corresponds to counter-clockwise rotation of the velocity vector, with the insect’s head viewed from the front (Fig. 1). Accordingly, when viewed from the mosquito’s head along the antenna the clockwise rotation corresponds to the increase in φ.

The resulting direction of the air vibration velocity in the stimulating system was determined by the vector superposition of the signals from both speakers. Changes in the sound wave direction relative to the mosquito in 15° (π/8) steps were accomplished by coordinated switching of voltage dividers in the SC transducer. For those angles at which the values of the functions sin(φ+45) or cos(φ+45) were negative, the signal polarity was inverted by switching the terminals of the speakers. This technique of variation of the sound wave vectors did not require any construction elements to be moved inside the test zone during the experiment, so it allowed to avoid vibrations which could affect the focal microelectrode recordings and, in addition, the measurements could be made faster compared to the techniques which involve the rotation of the speaker or the animal.

The moving parts of the speaker had a low resonant frequency (90 Hz). Due to the considerable response lag of the dome of the speaker and its support, the emission phase delay increased with the signal frequency up to the point of inversion. To stabilize the phase delay, a phase correction depending on the stimulation frequency was included in the speaker control circuit.

The sinusoidal stimuli were generated by the digital-to-analog converter LA-DACn10m1 (Rudnev-Shilyaev, Russian Federation). Acoustic calibration of the stimulating device was performed with an NR-231-58-000 differential capacitor microphone (Knowles Electronics, USA) attached to a micropositioner with axial rotation feature and set in the position of the mosquito. The same microphone put in 2 cm from the mosquito was used to record the stimulation signals during the recordings.

The differential microphone together with its amplifier was previously calibrated in the far field using the B&K 2253 sound level meter with a B&K 4135 microphone (Brüel & Kjær, Denmark). All sound level data in this study are given in the logarithmic scale in dB RMS SPVL (root mean square sound particle velocity level), with a reference level of 0 dB being equal to 4.85×10-5 mm/s, which corresponds in the far field to the standard reference sound pressure of 20 µPa.

### 2.3. Microelectrode recordings

Focal recordings from the axons of the antennal nerve were made with glass microelectrodes (1B100F–4, WPI Inc.) filled with 0.15 M sodium chloride and inserted at the scape-pedicel joint. In this study we preferred the extracellular responses to the quasi-intracellular ones due to the stability of the former over a long time interval required for directional measurements.

After the penetration of the cuticle electrodes had a resistance of 15–110 MΩ. The electrode was manipulated by means of micropositioner. The whole setup was mounted at the vibration-isolated steel table. Neuronal responses were amplified using a home-made DC amplifier (gain 10, input resistance >10 GΩ). To use the neuronal responses for feedback stimulation (see below) and to measure the response thresholds, the output of the DC amplifier was passed through an additional AC amplifier (gain 20, 30 or 40 dB, band-pass = 5–5000 Hz). Responses and stimulation signals were digitized using E14-440 A/D board (L-Card, Russian Federation) at 20 kHz sampling rate, and LGraph2 software. Digitized recordings were examined with Sound Forge Pro 10 (Sony).

Due to the fact that the electrode tip and the average diameter of sensory axon in the antennal nerve were of comparable size (1 μm or less), we cannot claim that the recordings were made from the individual axons. For the sake of simplicity, here we use the terms ‘unit’ or ‘sensory unit’ in the sense of one or several axons belonging to the primary sensory neurons of the JO, closely located within the antennal nerve and sharing similar frequency and phasic properties, thus representing a single functional unit. Detailed discussion of this issue can be found elsewhere (Lapshin and Vorontsov, 2013; Lapshin and Vorontsov, 2017)

### 2.4. Measurements of the directional sensitivity

While penetrating the antennal nerve by the electrode, the preparation was continuously stimulated with tonal pulses (filling frequency 200–260 Hz, amplitude 60-65 dB SVPL, duration 80 ms, period 600 ms). During this searching procedure the groups of the JO neurons situated orthogonal to the antenna oscillation could be overlooked. To avoid this, the vector of acoustic wave was periodically changed by 90° using the switch on the SC transducer.

The threshold measurements were made using either sinusoidal (to obtain absolute thresholds) or the positive feedback stimulation (to obtain relative thresholds). The essence of the latter method is a positive feedback loop established using the amplified in-phase response of a sensory unit as the signal to drive the stimulation loudspeaker. Applying such kind of stimulation to the sensory unit we expect it to ‘sing’ at a frequency which is close to its intrinsic tuning frequency – we call this effect ‘autoexcitation’. With feedback stimulation, the threshold was defined as the signal level which required one more incremental step at the attenuator output (+1 dB) for the system to enter the autoexcitation mode. With sinusoidal stimulation, the criterion of the response threshold was set at 2 dB of sustained excess of response amplitude above the average noise level in a given recording. At each combination of stimulation parameters the threshold was measured consequently at least twice.

To distinguish between the two above methods, hereinafter we will use the term ‘polar patterns’ for the results obtained by the positive feedback stimulation, and ‘directional characteristics’ for those measured with sinusoidal stimulation. It is important to bear in mind that the feedback method provides the unipolar response, i. e. it allows to distinguish between the two units responding to the opposite (180°) phases of the antenna vibration. The directional characteristics obtained using the sinusoidal stimulation are always bipolar.

The directional characteristics and polar patterns of sensory neurons were obtained by measuring the auditory thresholds at different angles of acoustic stimulation vector which was changed in 15° steps. Since the complete set of measurements took quite a long time (20-25 min), repeated measurements at certain angles (usually at 45° and 315°) were made not less than twice per measurement series to ascertain the stability of recording.

During the subsequent data processing, the maximum threshold value (Th _max_) was determined for a given recorded unit. Based on it, a set of derived values describing the unit directional characteristic or polar pattern was estimated by the formula A_i_ = Th _max_ − Th _i_. In the curves based on these data, the sectors of the highest sensitivity corresponded to the lowest recorded thresholds, and the central zero point corresponded to Th _max_. The angles at which no response at the best frequency was observed were given the value A_i_ = 0.

The angular sensitivity range of a unit was determined at –6 dB of the maximum sensitivity (in case of directional characteristics the two values from the symmetrical curves were averaged). The best angle of a given unit was determined as the bisector of this range.

Since it is known that frequency tuning of the JO, as well as the wingbeat frequency, highly depend on the ambient temperature (Costello, 1974; Villarreal et al., 2017), all the frequency data in this study underwent temperature correction to the value of 20°C. For such calculations, the previously estimated coefficient of 0.02 Hz per 1°C was used.

## 3. Results

### 3.1. Individual directional properties of the JO units

As a rule, in one specimen directional measurements were made consecutively from two or more recording sites within the antennal nerve. At each recording site the polar pattern of one unit (if a single unit was responding), two (in most cases) or three units was measured together with their tuning frequencies in the mode of feedback stimulation. Then, in case of a stable recording, the directional characteristics of the same unit(s) were measured using the sinusoidal stimulation. Recordings were made from 91 male mosquitoes. In total, directional properties of 306 units were measured in the frequency range 114–359 Hz, among them there were 46 single, 85 paired (two units recorded together but responding in antiphase) and 30 tripled units. In the latter group of recordings, two units responded in-phase, but demonstrated different frequency tuning, and one responded in antiphase and was tuned to the third frequency lying between the ones of the in-phase pair.

The examples of individual responses are shown in Fig.2. We do not show the waveforms here, they were similar to the ones descibed in detail elsewhere (Lapshin and Vorontsov, 2017).

**Fig. 2.**
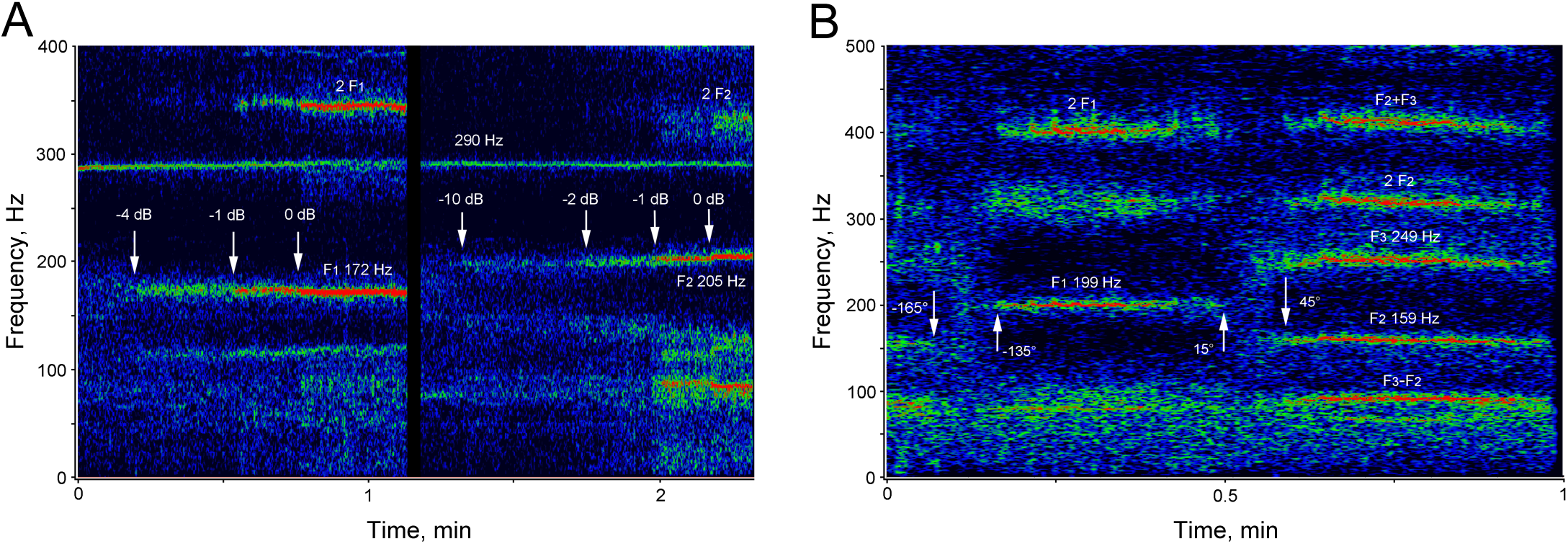
Examples of the JO unit responses to the feedback stimulation. A. The direction of sound wave is set to –60°, the feedback power is gradually increased from the sub-threshold levels, above –4 dB the response appears first as the higher level of noise, followed by sporadic bursts (from –1 to 0 dB) and continuous excitation at 172 Hz above 0 dB (absolute threshold of autoexcitation 42 dB SPVL at the fundamental frequency). Then, the stimulation is switched off, the direction of sound wave is rotated by 180° (to 120°) and the procedure repeated starting from –10 dB. The auto-excitation this time appeared at 205 Hz, threshold of autoexcitation 45 dB SPVL at the fundamental frequency. Continuous stripe at ca. 290 Hz represents the spontaneous activity in the JO, a correlate of active mechanics of the JO sensory cells, and it produces the combination harmonic (290–205=85 Hz), which can be seen in the right part of the sonogram. B. An example of triple unit system responding to the rotation of sound vector. The feedback level is kept constant 6 dB above the threshold of the first recorded unit (F1), the sound vector is rotated by 360° in 15° steps. First autoexcitation frequency, F1, appeared at –135° and disappeared at 15 (maximal level 52 dB at the fundamental frequency), then F2 (159 Hz, 50 dB) and F3 (249 Hz, 51 dB) appeared at 45° and disappeared at −165°. Note the combination (mixed) harmonics (F3–F2 and F2+F3) when two units were excited simultaneously. Arrows indicate the moments when the autoexcitation appeared and disappeared. Vertical axis: frequency, Hz, horizontal axis: time, minutes. Color represents the relative amplitude of response.

When the power of the feedback was increased from the sub-threshold levels, first a higher level of noise appeared (Fig.2A), followed by sporadic bursts of activity (from –1 to 0 dB in the shown example). At higher levels of feedback the response transformed into continuous excitation at the specific frequency, often with higher harmonics also present in the recording. When the direction of the sound wave was switched to the opposite, the kind of response was similar, but the unit(s) excited at the different frequency. We observed this effect in the previous studies when switching the phase of the signal in the feedback circuit, although here the change of the sound wave direction was done in a different way.

Alternatively, when the feedback power was kept stable above the threshold, and the vector of acoustic wave was rotated stepwise, the excitation gradually appeared and disappeared depending on the angle of stimulation. In triple unit recordings at certain angles two frequencies were present simultaneously, producing the combination, or mixed harmonics (Fig.2B).

In case of paired unit recording their individual polar patterns were opposite (180°±10°), mirroring each other, while their individual tuning frequencies were different (Fig3. A). For the same pair of units, the directional characteristic measured on the best frequency was bi-directional (figure-eight pattern) with its axis with slight deviation following the axis of previously measured combined polar patterns (Fig3. B). Angular orientation of polar patterns at the given recording site was arbitrary, with no obvious preference across the antennal nerve. Usually, after a slight axial shift of the electrode another pair or triplet of units started to respond, demonstrating different angular orientation and different tuning frequencies while maintaining the opposite, mirror-like polar patterns. The polar pattern of a single unit had the form of a petal located asymmetrically relative to the center of polar coordinates. One of the possible reasons for appearance of single-unit recordings may be the mechanical instability of the preparation due to the muscle contractions. In other words, some of the experiments ceased before the measurements from all directions were made. However, not all recordings with only a single unit responding can be explained in such a way.

**Fig. 3.**
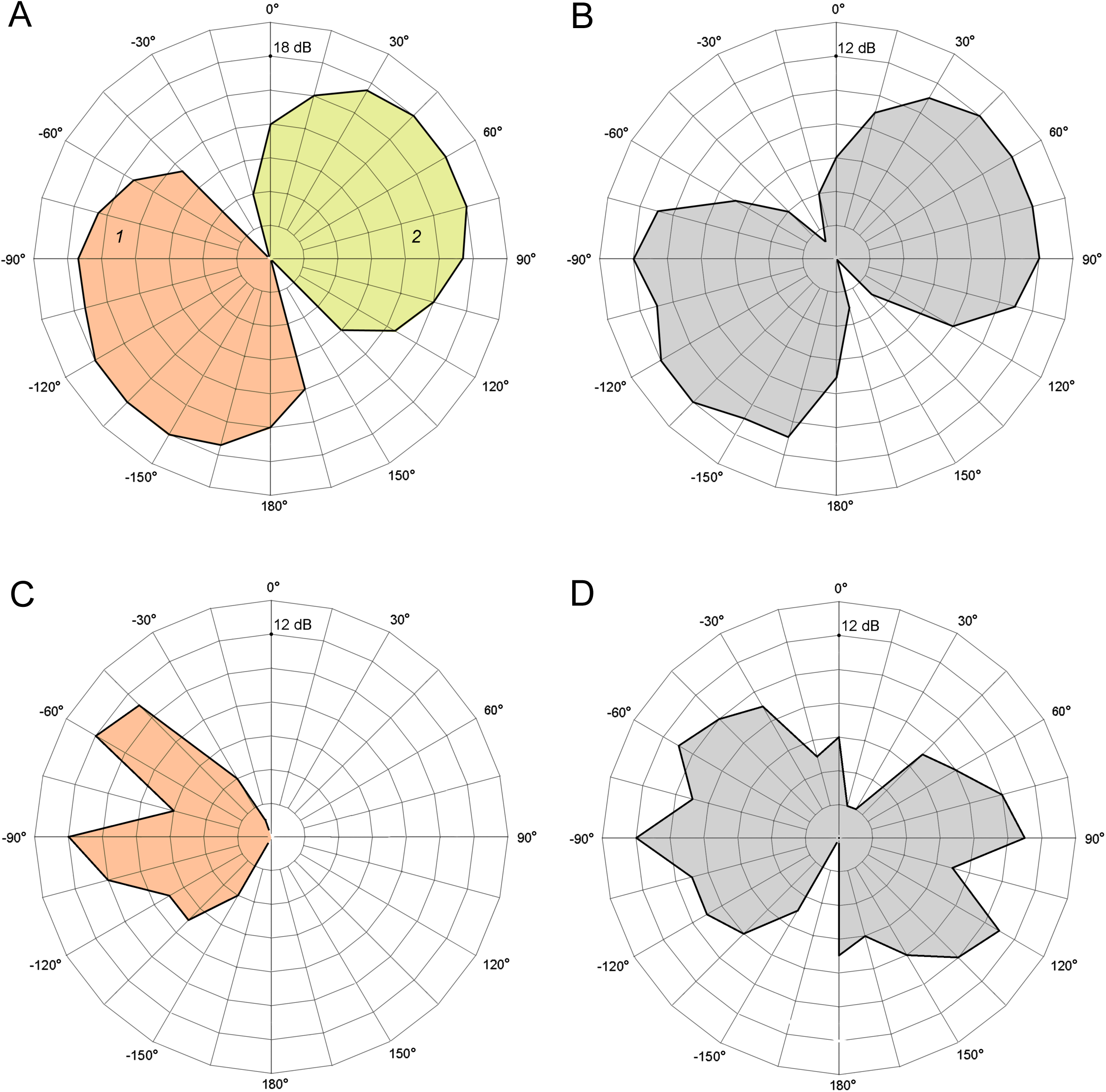
Examples of polar patterns and directional characteristics measured from the JO sensory units. A, polar patterns of a pair of antiphase units; best frequency of #1 is 201 Hz, of the #2 is 253 Hz. B, the same pair of units as in A, diagram obtained with 230 Hz sinusoidal stimulation, the threshold at the best direction is 29 dB SPVL. C, polar pattern of the single unit, best frequency at 199 Hz. D, the same unit as in C, directional diagram measured at 200 Hz, threshold 37 dB SPVL. Angle of sound wave is shown at the perimeter of each diagram, measured from the dorso-ventral axis (see Fig.1). Relative sensitivity is plotted radially in 3 dB (A) or 2 dB (B, C, D) steps.

Sometimes the ordinary petal shape of the polar pattern was distorted in the form of one or two notches appearing in it (Fig.3C). Directional characteristics measured from the same units demonstrated some similarity in shape (Fig.3D). Such distortions can be explained by the presence of additional antiphase units in the area of focal recording, and their effect on the recording quite predictably was more pronounced during the feedback stimulation and the polar pattern measurements.

The average angular sensitivity range of a unit, measured from directional characteristics at –6 dB from the maximum, was found to be 123° (σ=14.5°, n=74). The same, estimated from the polar patterns, was slightly narrower: 100° (σ=16°, n=275). This difference in estimates is easily explained, since the positive feedback, on which the measurements of the polar patterns were based, was very sensitive to the decrease in transfer coefficient, and this effect had to be most significant at the directions of minimal sensitivity of the unit.

The absolute thresholds of sensitivity at best frequencies varied from 22 to 44 dB SPVL (M=32 dB SPVL, σ=4.4 dB, n=74).

### 3.2. Directional characteristics of the JO

Both frequency tuning and individual sensitivity of units were found to be more or less evenly distributed around the antenna (Fig. 4). Some asymmetry in the angular distribution of units can be explained by the difference in the total numbers of units recorded in each sector (1 and 3 *vs* 2 and 4 in Fig. 4A). The absolute thresholds of sensitivity at best frequencies varied from 22 to 44 dB SPVL (M=32 dB SPVL, σ=4.4 dB, n=74).

**Fig. 4.**
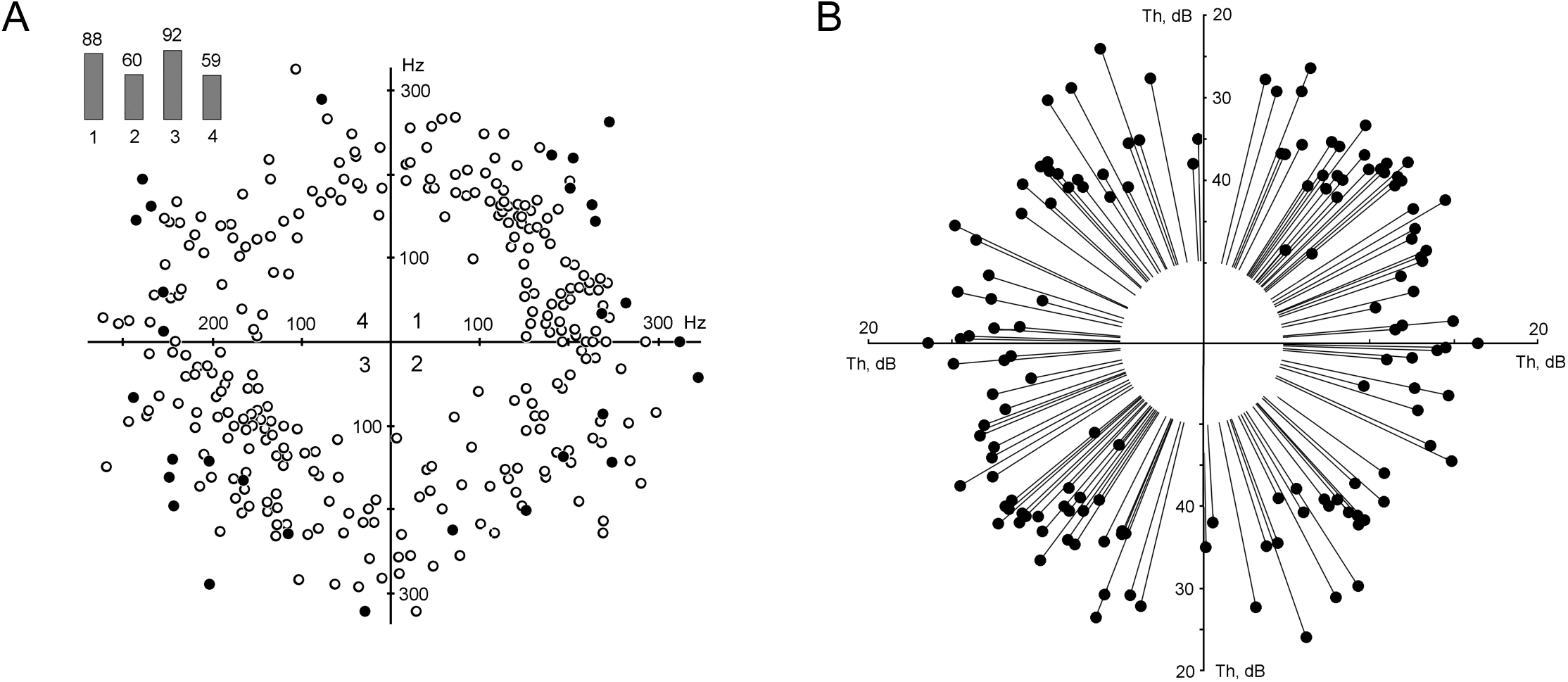
Directional properties of the JO units. All data are given in polar coordinates with the center corresponding to the axis of the antenna. A, radial distribution of best frequencies of the JO units. Measurements are made from polar patterns, obtained in the feedback stimulation mode, the individual tuning frequency values are plotted radially. Filled circles show the higher-frequency units (F3) belonging to the triple-unit systems. Histogram in the upper-left corner shows the total number of units recorded in each of the quadrants. B, radial distribution of individual thresholds. Measurements are made with sinusoidal stimulation at previously determined individual best frequencies, inverse thresholds are plotted radially, so the dots belonging to the higher sensitive units are further from the center of the graph.

### 3.3. Ratios between individual frequencies

The most remarkable feature demonstrated by the individual tuning frequencies of the JO units is the distinct relations between the ones belonging to the same recording site where a pair or a triplet of units responded simultaneously. We divided the whole dataset in two parts, corresponding to the pairs and triplets, and analyzed them independently.

The distribution of the ratio between the individual frequencies in the antiphase pairs (n=85) is shown in Fig. 5A. The major peaks are centered around 1.2 and 1.25. The integer representation of these values correspond to the ratios 6/5 and 5/4, respectively. The validity of representation of frequency ratio in the form of a fractional ratio of integers is supported by the values of the two minor peaks of the distribution: 1.17 ≈ 7/6 = 1.16(6), 1.3 ≈ 4/3 = 1.33(3).

**Fig. 5.**
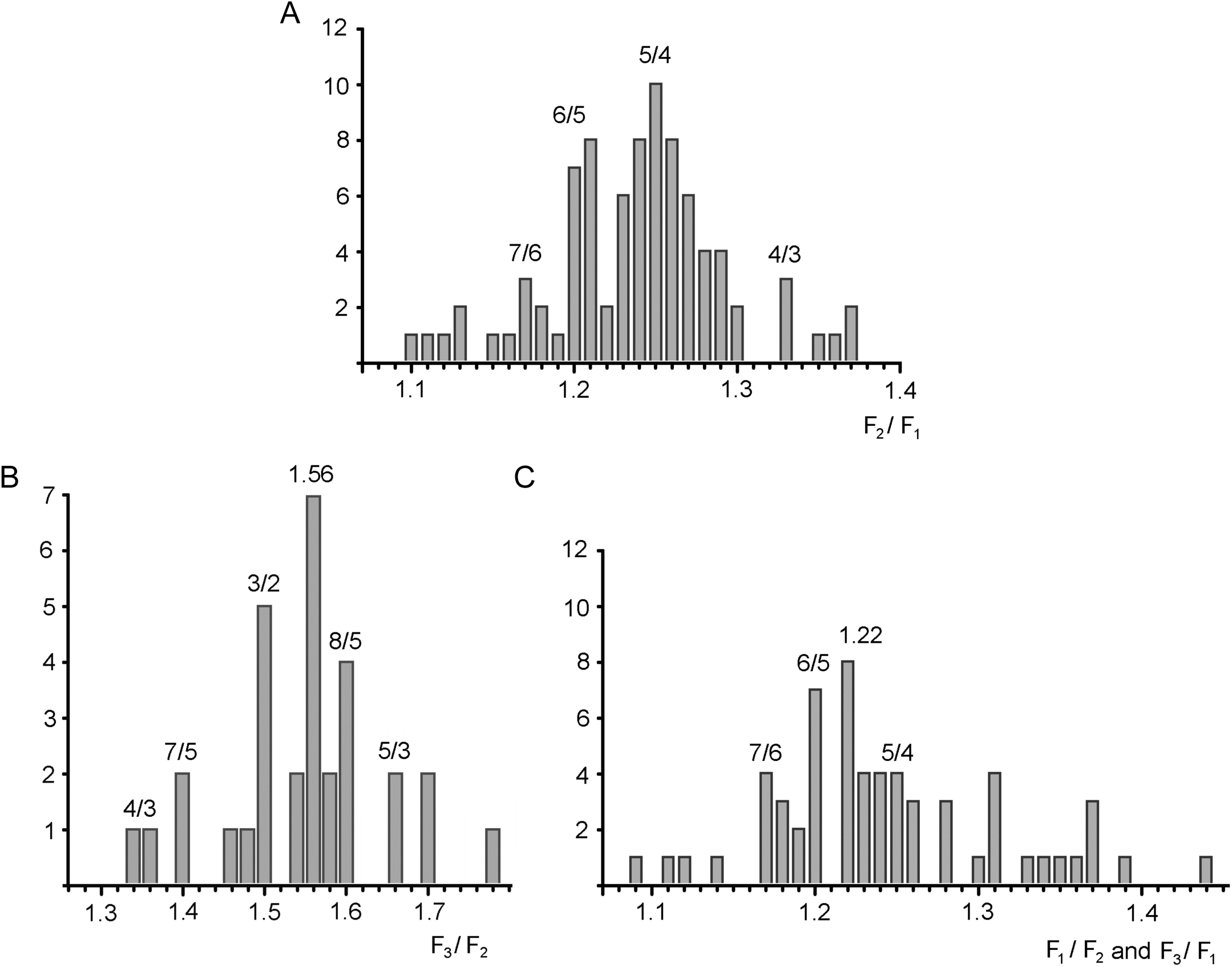
Distributions of frequency ratio between the units. Individual frequencies are designated as F1 and F2 for pairs and F1, F2 and F3 for triplets, where F1 unit is in antiphase to two in-phase units, lower (F2) and upper (F3) frequencies the example response of such is given in Fig.2B). Numbers above the histograms show the rounded values of the distribution local maxima and the respective integer ratios. A, pairs of antiphase units, n=85. B, C, triplets of units, n=30. B, ratios in the in-phase pairs. C, ratios in the antiphase pairs.

The scatter of actual ratio values can be easily explained by the accuracy of measurements in this study and the instability of frequency tuning in time: individual tuning frequencies were measured to 1 Hz, while during the measurements the frequency could spontaneously change within 2 Hz range. Both factors affecting the F1/F2 ratio could shift the resulting value by 0.01, or two bins of the histogram in Fig.5A, to either side of the mode. Within these limits, taken as the predicted variation, the ratio of 5/4 was characteristic of 38 pairs of antiphase units.

In triplets of units (n=30) two of them demonstrated the in-phase autoexcitation, meaning that each of the units received not only its own signal, converted to the vibration of the antenna, but also the one of the second unit, recorded simultaneously. Such cumulative effect led to the formation of a series of mixed harmonics in the recording. When the ratio between the frequencies was close to 1.5 (3/2, one of the peaks in the distribution in Fig.5B), the resulting spectrum became similar to a single harmonic series, like in Fig.2B. Such similarity complicated the identification of the individual tuning frequencies in the in-phase pair of units; however, the primary frequency could be distinguished from the mixed harmonics by the presence of the corresponding suppression zone at the same frequency, appearing after the rotation of the stimulation vector by 180°, thus converting the positive feedback to the negative one for a given unit (Lapshin and Vorontsov, 2017).

Figure 5B shows the distribution of frequency ratio F_3_/F_2_ from the triple systems (n=30). It has several peaks, the most pronounced at 1.56, while the one corresponding to the integer ratio 3/2 is lower. Similarly, in the distribution of the second frequency ratio from the same triplets (Fig. 5C) dominates the peak centered at 1.22 which does not correspond to some obvious integer ratio, but there are also peaks at 7/6 and 6/5 similar to the ones in Fig. 5A.

## 4. Discussion

### 4.1. Directional properties of the mosquito JO

The number of sensory units recorded in this study (n=306) represents only a minor part of their overall number in the JO. However, even this random sample of units, most probably, could perform the analysis of sound spectrum in all directions relative to the JO axis (Fig.4).

The average width of the directional characteristic is ca. 120° (Fig. 3B,D). Based on this estimate, we can conclude that 4-5 similarly tuned units, evenly distributed around the axis in the JO, would be enough to cover the whole directional range at the given frequency. However, this could be insufficient to provide the required accuracy of determining the angular coordinates of the sound source. According to Belton (1967) male mosquitoes are not attracted to the sounds which come from larger distance, even if these signals contain conspecific female-like tones. For small insects, the most accessible way to estimate the distance to a sound source is to measure its angular position during their own displacement in space (parallax estimation of distance). In calm weather, the swarming *C. p. pipiens* males fly longitudinal tacks. Analyzing the degree of parallax displacement of a sound source, they, apparently, can determine the distance to this source. One can presume that mosquitoes will pay attention to the sounds of nearby sources (within transverse size of the swarm) while more or less ignoring the sounds coming from larger distance. This would help to stabilize the position of the swarm and would increase the noise immunity of the male-female auditory communication channel. However, the task of instant triangulation demands high speed and precision of angular estimates performed by the JO, and can explain its seemingly redundant complexity.

Unfortunately, we cannot be sure that the diagrams of the Fig.4 show the true angular distribution of the differently tuned JO sensory units. The uncertainty lies in the possibility of selective recording from certain parts of the antennal nerve due to the geometric constraints in mutual arrangement of the mosquito, the recording electrode and the speaker. On the other hand, some asymmetry in the angular distribution of sensory units still must be present, since mosquitoes with one antenna maintained the ability, although much reduced, to locate a female (Roth, 1948).

### 4.2. Pairs and triplets of sensory units

Our measurements show that in the mosquito JO there is a large proportion or pairwise-combined units with different frequency tuning and oppositely oriented polar patterns: 85% of recorded units belonged to the paired or triple systems.

This finding means that during the single deflection of the antenna they generate antiphase electrical signals. It is attractive to speculate that this physiological finding corresponds to the well-known morphological fact that most sensillae in the JO contain two or three sensory cells (Boo and Richards, 1975a; Hart et al., 2011) and that their axons keep adjacent position further to the antennal nerve. The sinusoidal signals do not allow to separate responses of these two cells, but the positive feedback stimulation provides an opportunity to study antiphase units separately and measure their individual best frequencies and polar patterns.

How such antiphase responses are formed is currently unknown. We can propose at least two hypothetic mechanisms: one is based on the initially different polarity of the mechano-electrical transduction in the two cells belonging to a single sensilla, the other one – on the different and precisely adjusted latency of signal transduction in the two cells. The latter mechanism is, however, cannot work similarly in the wide range of frequencies. Currently we discard the third possibility – that antiphase axons belong to the units from the opposite parts of the JO capsule – since there is no morphological evidence of it.

The triple systems, including two units responding in-phase and the third one in antiphase, give additional insight to the underlying neuronal morphology and support the above speculations. Since in the triple-unit recordings the polar patterns of the in-phase pair were always oriented similarly, with high degree of confidence it can be assumed that the units producing these responses are morphologically combined in the capsule of the JO. Moreover, the specific ratio of best frequencies in such pairs and triplets indicates the functional interaction between these units (Fig.5).

Morphological combination of two or more sensory cells into the sensilla is known for many insect chordotonal organs (Field and Matheson, 1998), including the one of Drosophila flies, Chironomidae midges and mosquitoes. From the widespread occurrence of this phenomenon among insects, one can assume that it must have some general functional significance, not specific to the mosquitoes. One of the possible tasks performed by such organization of the sensillae may be preventing the auditory neurons from sending to the brain the responses to large low-frequency deflections of the antenna caused by wind currents during the flight maneuvers of an insect. The antiphase pair of sensory cells, having equal sensitivity in low-frequency range, can filter out such signals even before they leave the JO or the antennal nerve, provided that these cells are interconnected by gap junctions. The latter was indeed demonstrated in the JO of Drosophila (Sivan-Loukianova and Eberl, 2005). Such mechanism must be very sensitive to the similarity of parameters of both sensory cells. Combining them into a single sensilla is fully justified in order to ensure equality of their directional characteristics and similarity of metabolism.

In a pairwise combination of specifically tuned antiphase units one can notice an analogy with the opponent coding of color information in the vertebrate retina (Daw, 1973). The opponency of auditory sensory units with different frequency tuning can substantially facilitate the following information processing in the brain since it allows to easily distinguish the sounds with continuous (noise-like) spectrum from the ones with line spectrum like the sound of a flying female, or to produce selectivity for other stimulus features (Chang et al., 2016).

### 4.3. Ratios between the individual frequencies

The tendency of units in paired and triple systems to have their best frequency ratios equal to the simple integer fractions (Fig.5) may be a sign of the mechanism of signal processing, some kind of internal ‘language’ of the system, representing the auditory space of mosquito. Remarkably, almost similar frequency ratios between the paired flight tones of male and female mosquitoes were observed (Aldersley et al., 2016). Such tendency may also explain the multi-modal shape of overall distribution of individual frequencies, which was demonstrated in our previous study (Lapshin and Vorontsov, 2017).

It should be noted that in triple-unit systems the ratios between the individual frequencies are interdependent. For example, if the ratio in the in-phase pair F3 / F2 = 1.5 and the ratio in any of the antiphase pairs from the same triplet is, for example, F3 / F1 = 1.25, then the ratio in the other antiphase pair should be equal to F1 / F2 = 1.2 (F1 / F2 = F3 / F2 : F3 / F1). However, there is a possible alternative, when in triple systems the primary ones are the ratios in the antiphase pairs. For example, if F1 / F2 = 5/4 and F3 / F1 = 5/4, then the third ratio between the in-phase units becomes dependent: F3 / F2 = 25/16 = 1.5625. Similarly, with the two other frequency ratios characteristic of the antiphase pairs equal to F1 / F2 = 4/3 = 1.33(3) and F3 / F1 = 7/6 = 1.166(6), for the dependent in-phase pair we get F3 / F2 = 14/9 = 1.55(5). Remarkably, in the Fig.5B the above calculated ratios fall into the major peak of the distribution. Apparently, the frequency ratios between the individual units in the triple systems do not follow clearly defined criteria (as it seems they do in pairs of units, Fig. 5A), at the same time being distinctly non-random. Most probably, they are limited by the more strict conditions which simultaneously connect three elements, for example F3 / F2 × F3 / F1 = 1.22 × 1.22 = 1.5, or 1.22 × 1.28 = 1.56. Sub-peaks centered at these values are indeed present in the distribution (Fig.5B,C).

The seeming complexity of frequency ratios in the JO may be explained in the framework of primary signal processing. Since every sound, even the pure tone, comes to the JO sensory units of a flying mosquito accompanied by mixed harmonics, there must be a demand to analyze this complex auditory image and to simplify the input to the brain interneurons. Highly parallel system of the JO sensory units supplemented by the ability to instantly discriminate the certain combinations of tones seems to be almost perfectly suited for the task.

However, the complex pattern of frequency ratios that we observed could simply originate from the known phenomenon of ′harmonic synchronization′, that is the specific mode of interaction between the coupled resonant nonlinear systems when their frequencies are integrally related to each other. The stability of such synchronization is determined by the local decrease in the energy of the entire system (Yang et al., 2012) and generally decreases with increasing frequency multiplicity. One cannot exclude the hypothesis that the specific frequency ratios, including the ones which manifest themselves in behavior, are not a part of signal processing mechanism of the JO, but just a by-product of energy optimization.

However, regardless of the functional meaning of harmonic synchronization, it is possible only if the oscillators express spontaneous activity at their best frequencies and interact, mechanically or electrically. In the JO sensory cells the good candidate mechanism of interaction would be the active auditory mechanics (Göpfert and Robert, 2001) based on the dynein – tubulin motor of the ciliated sensillae (Warren et al., 2010). Such kind of interaction can also explain the appearance of mixed harmonics visible in the sonograms of Fig2A.

Our recent finding suggests that mosquitoes can potentially demonstrate different kinds of responses to different frequencies of sound. It was shown in behavioral tests that *Aedes diantaeus* mosquitoes demonstrate fast avoidance response in the frequency range 140-200 Hz (Lapshin and Vorontsov, 2018). In these experiments mosquitoes which were previously attracted by the sound imitating the wingbeat tone of a female (280–320 Hz) left the stimulation area within one second from the onset of the test signal (amplitude 57–69 dB SPVL), flying up, sideways and backward relative to the direction of test signal arrival. These and other behavioral observations, together with our current physiological findings, strongly suggest that the JO of mosquitoes can discriminate tones coming from different directions in a wide range of amplitudes.

## 5. Acknowledgments

The field facilities for this study, Kropotovo biological station, was provided by the Koltzov Institute of Developmental Biology RAS.

